# Persistence of SARS-CoV-2 virus and viral RNA on hydrophobic and hydrophilic surfaces and investigating contamination concentration

**DOI:** 10.1101/2021.03.11.435056

**Authors:** Susan Paton, Antony Spencer, Isobel Garratt, Katy-Anne Thompson, Ikshitaa Dinesh, Paz Aranega-Bou, David Stevenson, Simon Clark, Jake Dunning, Allan Bennett, Thomas Pottage

## Abstract

The transmission of SARS-CoV-2 is likely to occur through a number of routes, including contact with contaminated surfaces. Many studies have used RT-PCR analysis to detect SARS-CoV-2 RNA on surfaces but seldom has viable virus been detected. This paper investigates the viability over time of SARS-CoV-2 dried onto a range of materials and compares viability of the virus to RNA copies recovered, and whether virus viability is concentration dependant.

Viable virus persisted for the longest time on surgical mask material and stainless steel with a 99.9% reduction in viability by 124 and 113 hours respectively. Viability of SARS-CoV-2 reduced the fastest on a polyester shirt, with a 99.9% reduction within 2.5 hours. Viability on cotton was reduced second fastest, with 99.9% reduction in 72 hours. RNA on all the surfaces exhibited a one log reduction in genome copy recovery over 21 days.

The findings show that SARS-CoV-2 is most stable on non-porous hydrophobic surfaces. RNA is highly stable when dried on surfaces with only one log reduction in recovery over three weeks. In comparison, SARS-CoV-2 viability reduced more rapidly, but this loss in viability was found to be independent of starting concentration. Expected levels of SARS-CoV-2 viable environmental surface contamination would lead to undetectable levels within two days. Therefore, when RNA is detected on surfaces it does not directly indicate presence of viable virus even at high CT values.

**Importance:** This study shows the impact of material type on the viability of SARS-CoV-2 on surfaces. It demonstrates that the decay rate of viable SARS-CoV-2 is independent of starting concentration. However, RNA shows high stability on surfaces over extended periods. This has implications for interpretation of surface sampling results using RT-PCR to determine the possibility of viable virus from a surface. Unless sampled immediately after contamination it is difficult to align RNA copy numbers to quantity of viable virus on a surface.

## Introduction

Severe acute respiratory syndrome coronavirus 2 (SARS-CoV-2) causing coronavirus disease 2019 (COVID-19) has spread globally and many countries are experiencing ongoing local transmission despite varying levels of control efforts. SARS-CoV-2 is primarily transmitted via respiratory droplets from an infected host (1). Studies have confirmed aerosol viral transmission (2–4) with SARS-CoV-2 being shown to remain viable in aerosols for between 90 minutes and 3 hours in laboratory studies (5, 6). Infections from direct person to person transmission have been confirmed as well as indirect transmission through close contacts after tracing of case clusters (7, 8). It is suspected that contaminated surfaces or fomites may also have a role in transmission. Studies detailing SARS-CoV-2 viability on surfaces have contributed to this (6).

SARS-CoV-2 RNA has been detected on environmental surfaces, potentially indicating the presence of the viable virus (9, 10). Current environmental sampling of surfaces using swabs, primarily uses real-time PCR (RT-PCR) to detect viral genome in samples. Few studies have been able to isolate viable virus from environmental surface sampling, even where RT-PCR indicates a high level of SARS-CoV-2 RNA is present (11). Recent manuscripts have identified survivability ranges for SARS-CoV-2 on surfaces in the laboratory, with only one demonstrating the relationship between viable recovered virus and RNA on the surface (12, 13). As the risk of infection from of virus-contaminated surfaces is difficult to predict (14), further investigation is required to enhance understanding on the survivability of SARS-CoV-2 on surfaces.

The three aims of this study were; (i) measure the persistence of viable SARS-CoV-2 virus on common personal protective equipment (PPE) materials (both hospital-grade and reusable fabrics), high-touch surface materials and commonly worn fabrics; (ii) to investigate the relationship between recoverable viable virus from these surfaces and the levels of SARS-CoV-2 RNA in the same sample; (iii) determine the relationship between inactivation rate and initial viral titre load on surfaces.

## Results

SARS-CoV-2 viability decreased on all materials during the 2.5 hour drying period, on average by 1.01 log_10_ (between −0.18 log_10_ for disposable gown to −3.66 log_10_ for polyester sports shirt, standard deviation 1.06 log_10_) from a high starting inoculum of approximately 4 × 10^5^ pfu per material. Viable virus could be recovered from the surgical mask and stainless steel coupons for the longest periods of time (log_10_ reductions of 4.91 and 4.99, respectively, over 7 days). The recovery time for cotton t-shirt and polyester sports shirt materials was shorter (log_10_ reductions of 5.15 over 5 days and 3.9 within 1 day). RNA copy number was recovered at higher concentrations in all samples compared to the levels of viable virus, decreasing by ~1.5 log_10_ (non-porous, hydrophobic) and ~1 log_10_ (porous, hydrophilic) over the initial 7 days, then stabilised at around 10^7.5^ copies per coupon from day 7. The ratio of viable virus recovered ranged from 10^3^ to 10^8^ times less when compared to viral RNA assayed, from start to finish of the study period (Fig 1).

**Fig 1.**
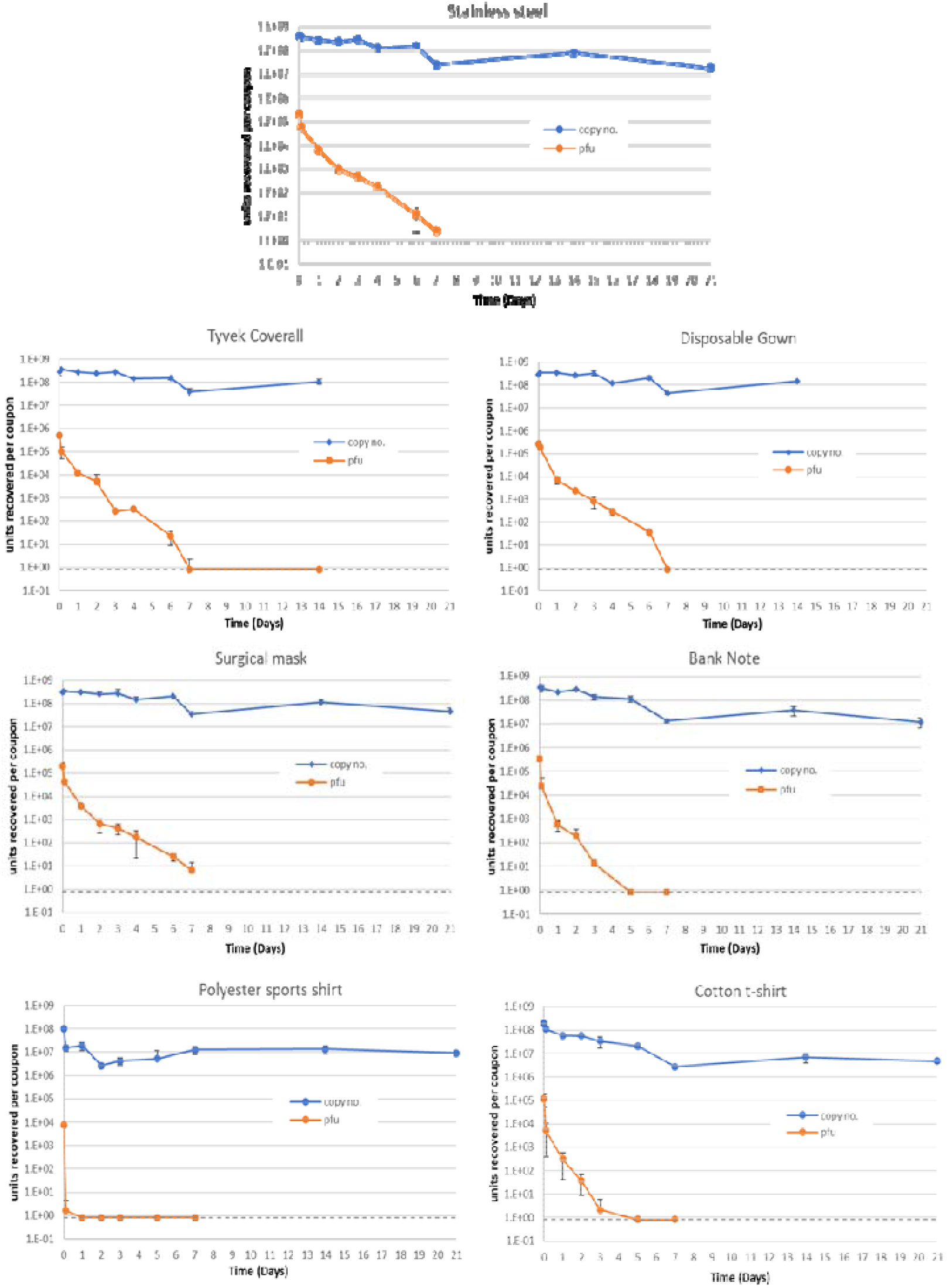
Mean quantities of viable virus recovered (pfu/coupon, orange) and viral RNA detected (genome copy number/coupon, blue), for 7 materials assessed. Error bars represent the standard deviation from three replicates. Grey dashed line represents the limit of detection of the plaque assay for the combined assays from the triplicate coupons (0.8 pfu/ml). For Tyvek coverall and disposable gown, 21 day coupons were not processed.

Linear regression analysis was completed using the recoverable virus data to calculate time for percentage reduction (Table 2). Regressions were calculated from t=2.5 hours onwards. Calculated decay rate is fastest on cotton with a 99.9% reduction in recovery within 72 hours. The longest survival of virus is observed on surgical mask material, where 124 hours is required for a reduction of 99.9%. For the polyester sports shirt a >3 log_10_ reduction was detected during the initial 2.5 hour timepoint (Table 2).

**Table 1.** Materials used in this study.

| Material | Properties |
| --- | --- |
| Stainless steel, Tyvek, disposable gown, bank note | Hydrophobic, non-porous |
| Surgical mask | Hydrophobic, porous |
| Cotton T-shirt, polyester sports shirt | Hydrophilic, porous |

**Table 2.** Time for percentage reduction values for the multi-surface study. Data is sorted in descending order by time to 99.9% survival rates. N/A = not applicable.

| Material | Log change<br>after 2.5 hour<br>drying | Time for percentage reduction (hours) |  |  |  |
| --- | --- | --- | --- | --- | --- |
|  |  | R<sup>2</sup><br>value | 90% | 99% | 99.9% |
| Surgical mask | -0.64 | 0.954 | 27.9 | 76.2 | 124.4 |
| Stainless steel | -0.56 | 0.982 | 30.7 | 71.6 | 112.6 |
| Tyvek Coverall | -0.70 | 0.962 | 33.3 | 69.3 | 105.2 |
| Disposable Gown | -0.18 | 0.946 | 23.0 | 59.7 | 96.5 |
| Bank note | -1.13 | 0.956 | 17.5 | 45.1 | 72.7 |
| Cotton t-shirt | -1.34 | 0.904 | 17.2 | 44.6 | 72.0 |
| Polyester sports shirt | -3.66 | N/A |  | <2.5 |  |

The results from the comparative study involving two viral titres revealed an initial rapid decrease in the recovery of viable virus from the surface during the drying period, with the inactivation rate decreasing after drying. Virus was recovered after four days for the low inoculum and up to seven days for the high inoculum (Figure 2). Parallel survival rates of high and low inocula demonstrate that the decay rate of SARS-CoV-2 is independent of concentration when applied to a stainless steel surface (P>0.05).

**Fig 2.**
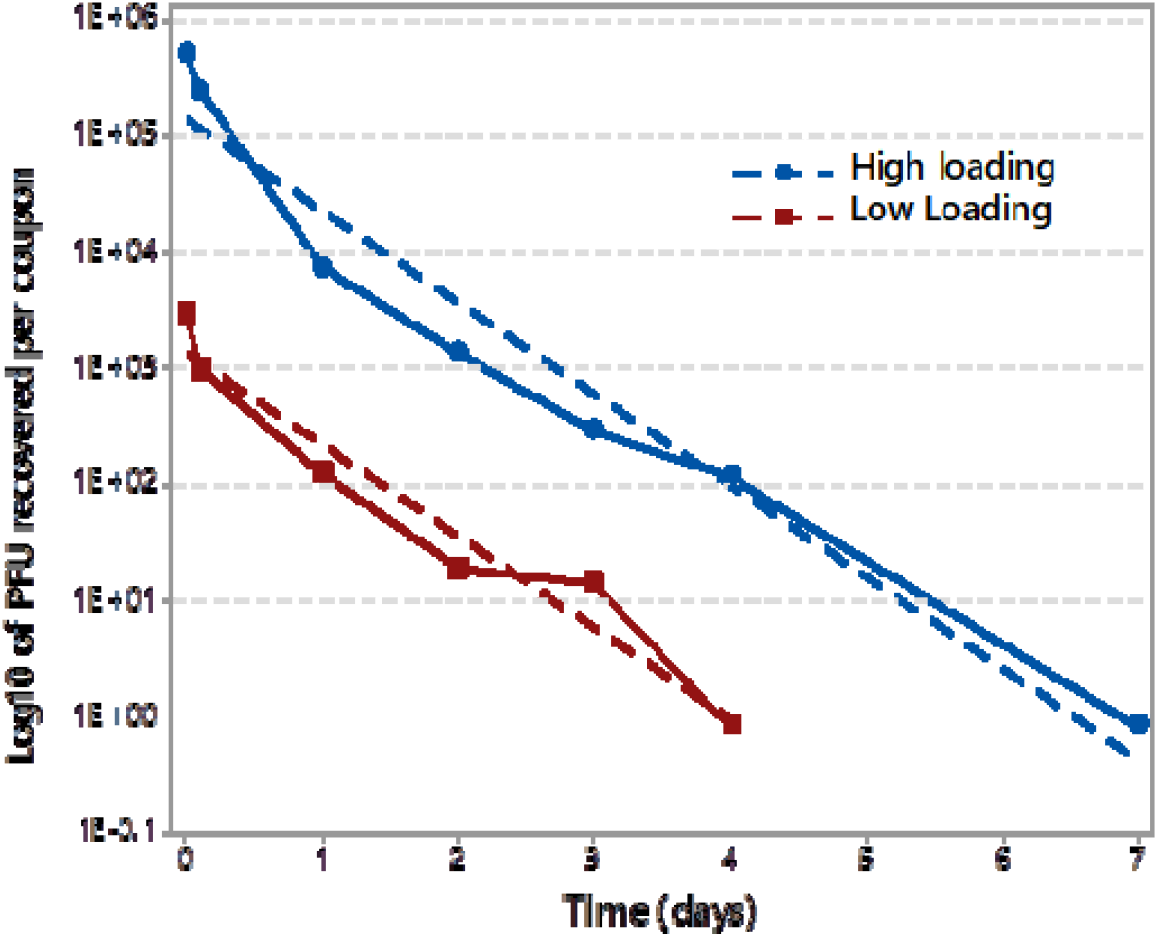
Virus viability results from loading of high (4 × 10^5^ pfu added, blue) and low (4 × 10^3^ pfu added, red) SARS-CoV-2 inoculum onto stainless steel coupons (n = 3). Dashed lines show linear regression based on recovery over time.

## Discussion

Contact with SARS-CoV-2 contaminated surfaces is thought to be a route of transmission in the current pandemic (8, 15). Surfaces can be contaminated by virus-containing droplets generated from an infected individual or contact with contaminated hands, with potential onwards transmission via direct surface contact (16). With contamination events likely to occur in a range of materials, this study investigated the survival of SARS-CoV-2 UK isolate hCoV-19/England/2/2020 and associated viral RNA on a range of surfaces that are at risk from droplet and touch contamination.

Existing studies examining time-based viability of SARS-CoV-2 on different surfaces have focused on a single virus titre (6, 12, 13, 17). Our work investigated the persistence of high titres of SARS-CoV-2 on various surfaces, and at two different titres on stainless steel coupons with identical conditions.

When inoculated onto stainless steel, a five-log reduction in viability of the UK SARS-CoV-2 isolate England 02/2020 HCM/V/052 was observed over a 7 day period (Figure 1). Riddell *et al* showed a similar log_10_ reduction at 28 days and Kasloff at 14 days, using the Australian isolate Betacoronavirus/Australia/SA01/2020 and Canadian isolate hCoV-19/Canada/ON-VIDO-01/2020 respectively (12, 17). In both studies, the viral propagate included additives such as serum and mucin to mimic bodily secretions (12, 17). Where in this current study the virus stock suspension was centrifuged to remove the majority of the cell debris, but left salts and proteins from the growth media used to propagate the virus. The initial starting inoculum concentration (between 10^5^−10^6^ pfu per surface) and environmental conditions of temperature and RH are similar for the two previous studies and this study, demonstrating it is likely that the differences observed in survival results could primarily arise from a protective effect afforded by added serum/mucin and/or the different isolates used. Whilst potentially artificially high, the starting inoculum in this current study provides the ability to determine the inactivation characteristics of the virus on the different materials which a lower starting inoculum may not.

Currently there are no published studies investigating inter-isolate differences in environmental surface stability of SARS-CoV-2. A study investigating the stability of SARS-CoV-1 (AY274119.3) and SARS-CoV-2 (nCoV-WA1-2020) by van Doremalen et al showed that both were similar under their experimental test conditions (6). Chin et al have reported similar findings to the work presented in our study using a comparable starting inoculum, also without additional protein (13). Their results showed that infectious SARS-CoV-2 was recovered from a banknote and stainless steel on days 4 and 7 respectively, compared to our results of recoverable virus on day 5 from the banknote and recovery on day 7 from stainless steel. With little evidence of difference in environmental stability between isolates of SARS-CoV-2 the addition of bovine serum albumin and mucin to the inoculating suspensions indicates that additional protein provided a protective effect to the virus during and after drying onto the surfaces (12, 13, 17).

Droplets on non-porous or hydrophobic surfaces dry in a beaded shape, giving a high volume to surface area ratio. In such an environment, these droplets can produce a core-shell structure (18), which can concentrate the virus particles, salts and organic material into smaller clumps (19). Such clumping is seen often in nature due to association with cellular matter or protein (20, 21). These closely associated virus particles are protected from environmental pressures such as desiccation, UV and heat, which cause inactivation (20). However, on porous or hydrophilic surfaces the droplets are absorbed into the material across a larger surface area, which will lead to less clumping and to the presentation of individual viral particles; this may confer less protection from the drying effects of the environment, leading to a reduction in the viability of the virus.

Whilst other studies have designated materials as porous and non-porous, this may be an oversimplification of the surfaces studied (12, 17). The surface of a surgical mask is porous but is made up of overlapping hydrophobic fibres; similarly, Tyvek material is produced with non-woven fibres of high density hydrophobic polyethylene, but presents microscopic pores on the surface. In the context of our study, relatively low amounts of liquid are being added to these surfaces. Thus, these small droplets of liquid cannot penetrate into the materials, as their hydrophobicity ensures the droplets of liquids remain on the surface of the material during the drying process; making the surfaces behave more like a non-porous one. Our results show that the porous but hydrophobic surfaces of the surgical mask, disposable gown and Tyvek coverall produce similar decay rates when compared to the non-porous hydrophobic surfaces of stainless steel with a five log_10_ reduction in recovered infectivity over 7 days. Viable SARS-CoV-2 was recovered from these surface materials over longer periods of time compared to the truly porous and hydrophilic surfaces tested, cotton and woven polyester. An exception was the hydrophobic polymer bank note, from which viable virus was recovered at the limit of detection for days 5 and 7 of the study. This is different to the study of Riddell et al where the recovery from the Australian bank note was similar to the other non-porous surfaces tested (17). At present it is not clear why this surface had decreased viability compared to the other non-porous hydrophilic surface used in this study, although there may be antiviral properties from some of the dyes used in the bank note.

Following a 4.73 log_10_ decrease, infectious virus was recovered from cotton material up to 3 days after inoculation; matching previous studies, reporting more rapid inactivation of virus particles on cotton surfaces compared to others (12, 17). These results may be attributed to two factors unrelated to any potential anti-viral activity of the material: retention of virus within the cotton fibre matrix, or losses during the inoculum application due to wicking. Due to the cotton's hydrophilic, woven nature, the liquid inoculum rapidly absorbs and penetrates into the fibres which, when dried, might cause interactive forces, limiting the release of virus particles, which is shown by a greater than 1 log_10_ reduction in recovery of viable virus after the drying period. This decrease in detection of viable virus may therefore be attributable to inefficient recovery from this specific type of material rather than increased inactivation. This result indicates that viral particles may remain in cotton fibres after contamination posing a forward transmission risk, but they will likely not be released from the substrate to cause infection. To counteract the materials inherent absorbent nature, during the inoculation and drying steps we suspended the cotton in strips across an open box. Whilst this exposed the virus inoculated coupon to the environmental conditions on both sides of the coupon it reduced any potential losses of the virus due to wicking on to container surface from the coupon as seen in a previous study (12).

Though polyester, produced from polyethylene terephthalate, is hydrophobic, when spun into fine fibres, aligned in the same orientation and woven into fabric, it behaved like the other woven fabric tested, cotton. It is possible that the aligned polyester fibres which are close together, but not fused, causes capillary action to draw the liquid into the interstitial spaces between the fibres and trap the virus particles. Virus that was inoculated onto polyester sports shirt was rapidly inactivated to unrecoverable levels in one day; there may also be interaction between the chemicals used to process/colour the fabric and the virus (22).

SARS-CoV-2 RNA has been detected on surfaces in different environments, but there have been few reports of viable virus recovery from these surfaces. The use of RT-PCR to determine the presence of SARS-CoV-2 on surfaces has advantages, increased sensitivity (RT-PCR can detect small amounts of target RNA) and rapid high throughput of samples compared to culture-based methods. The limitation of the use of RT-PCR in such studies is its inability to distinguish between viable and non-viable virus.

The detection of SARS-CoV-2 RNA from surface samples is used to indicate that virus (viable or non-viable) has been present on that surface at some point previously. Lower cycle threshold (CT) values from the RT-PCR assay indicates that more copy numbers of target RNA are present in that sample. Our study determined that a CT value of 18 equates to approximately 5 × 10^8^ copy numbers of the RNA target. The initial recovery of infectious virus from the materials (Figure 1), excluding the polyester sports shirt, is approximately 3.1 log_10_ (SD 0.13 log_10_) lower than the copy number, showing that there is a large amount of RNA exogenous to infection-competent viral particles in the inoculum. This difference between infectious virus and RNA copy number was also reported by Kasloff et al (12). The ratio between copy number and viable virus changes with time with recoverable infectious virus rates reducing more rapidly than copy numbers recovered in the same sample over time. The RNA is therefore more persistent in the environment compared with infectious virus. It has been shown that SARS-CoV-2 RNA was detected on cruise ship surfaces 17 days after cabins had been vacated (23).

Thus, it is not possible to draw conclusions on the viability of surface contamination from genome copy number of RNA detected after the initial contaminating event. In addition, the comparative persistence of RNA on the surfaces in comparison to infectivity makes it difficult to relate copy number to the date when the contamination may have occurred. Although our laboratory-based study used a concentration of infectious virus that may not reflect the contamination load present in the environment, we demonstrated recovery of viable virus at high and low concentrations.

Patient nasal and throat swab samples have produced CT values below 18 (24, 25), even to a CT value of 10 (26), but reported surface samples have produced CT values above 28 (9, 27–29). Using the results from our study to provide a calculation of the initial viable load on the surface, a CT of 28 would provide an approximate infectious virus titre of 10^2^ viral particles. Using the recovery results from stainless steel as a representative surface in this study (Figure 2), infectious virus from an initial recoverable inoculum of 10^2^ viral particles would be unrecoverable within 2 days. This is based on the vortex mixing recovery method used and the detection would be thought to reduce further using direct surface sampling methods. Future studies could address this limitation of knowledge where the copy number is determined for much lower concentrations of SARS-CoV-2 on surfaces which will help to further identify if the persistence of RNA is independent of concentration and addressing the relationship to viable virus recovery.

## Conclusions

This study shows that the UK SARS-CoV-2 isolate, hCoV-19/England/2/2020, remains viable on hydrophobic surfaces for up to 7 days, with recoverable viability on hydrophilic surfaces reduced to 3 days at ambient temperature and relative humidity. Indicating that some common surfaces could pose an infection risk if contaminated with high concentrations of virus, although viable virus contamination levels of environmental surfaces are likely to be at a low concentration. In contrast, recovery of RNA from the same samples shows little reduction in copy numbers over the same period. The data presented also indicates that the inactivation rate on environmental surfaces is independent of initial loading for SARS-CoV-2 and varies depending on surface type.

## Materials and methods

### Viral isolate

The SARS-CoV-2 isolate, England 02/2020 (EPI**_**ISL**_**407073) passage 3 (P3) used in this study was propagated by the High Containment Microbiology Department at PHE, Porton Down. The virus was isolated from a clinical sample taken during acute phase illness, using Vero E6 cells (ECACC, 85020206). A P2 master bank was produced using Vero E6 cells (BEI Resources, NR-596) and a P3 working bank produced in Vero/hSLAM cells (ECACC 04091501). Cell lines were infected at 95% confluence with an of MOI 0.0005 – 0.01 and maintained in 1x Minimal Essential Medium GlutaMax, 4% heat-treated foetal bovine serum (Gibco), 1x non-essential amino acids (Gibco), 25mM HEPES buffer (Gibco); additionally, Vero/hSLAM cells were also maintained in the presence of 0.4mg/ml geneticin (ThermoFisher Scientific, Gibco). Virus was harvested 3-6 days post-infection and supernatant clarified by centrifugation (3000 rpm, 10 min), virus was aliquoted and stored at −80oC. The titre of the P3 virus stock was determined to be 2.0 × 107 pfu/ml by plaque assay. All work handling SARS-CoV-2 was performed within a containment level 3 laboratory.

### Preparation of test surfaces

The surfaces used in this study are representative of non-porous hand-touch sites (stainless steel, 316 grade) and bank note (English polymer £10 note); PPE items used in the hospital and wider environments (Multiple layered surgical mask, Tyvek coverall, disposable plastic gown (37310 Breathable Impervious Gown)); and materials representing clothing items; cotton t-shirt (Fruit of the Loom) and polyester sports shirt (85% polyester, 15% elastane, Activewear SFP5-M02). 1cm × 1cm coupons of each material were prepared. Once prepared, non-porous coupons (stainless steel and bank note) were cleaned with Neutracon (NEU5, Scientific Laboratory Supplies, Nottingham, UK) detergent followed by rinsing with 70% IPA. Additionally, stainless steel was sterilised by autoclaving. The surfaces not suitable for cleaning (disposable plastic gown, cotton t-shirt, polyester sports shirt and Tyvek coverall) were purchased new and handled aseptically. Materials were subdivided depending on their surface properties, Table 1.

### Inoculation of test surfaces

Viral aliquots were thawed to room temperature immediately prior to inoculation of the materials. The stock suspension was diluted with cMEM to give a starting concentration of 4 × 10^7^ and 4 × 10^5^ pfu/ml for high and low loading, respectively. Coupons were inoculated by the addition of 2 × 10μl droplets of virus culture within a negative pressure flexible film isolator (FFI) and were left uncovered in plastic petri dishes in the FFI for the duration of the study at a temperature of 21.5°C (±1°C) and an average relative humidity of 45%. Three biological replicate coupons were prepared and inoculated for each time point (hours; 0, 2.5, 24, 48, 72, 96, 144, 168, 336 and 504) and each material. Triplicate coupons exposed but not inoculated with the virus acted as negative controls.

### Recovery of SARS-CoV-2 from test surfaces

Recovery of virus from the coupons at each specified timepoint was performed by transferring a coupon into 1ml of cMEM in a 7ml bijou tube with 4 glass beads (3mm diameter), followed by vortex mixing for 1 minute at maximum (Heidolph Multireax vortex). The resulting suspension was transferred to a cryotube for storage at −80°C before analysis. Storage of samples at −80°C was not found to affect viability of the virus or denature the RNA (results not shown). Timepoint zero (t=0) coupons were recovered within 5 minutes of being inoculated, before any drying occurred.

### Plaque Assay

Coupon recovery liquid was assayed by thawing samples at room temperature, then serially diluted (1 in 10) with MEM ((Gibco), 1% L-glutamine (Gibco), 1% Non-Essential Amino Acids and 2.5% HEPES 1M). 100μl of each dilution was pipetted in duplicate (technical replicates), up to four replicates for neat dilutions (400μ) onto confluent vero-E6 cells within a 24 well plate (7.9 × 10^4^ cells/cm^2^). After one hour of incubation (±15 minutes) at 37°C with plate rocking every 15 to 20 minutes, 0.5mL CMC overlay was added to each well; (1.5% CMC (3% (w/v) carboxymethylcellulose solution in sterile distilled water (Sigma C4888)), 1% Antibiotic Antimycotic Solution (100x) (Sigma Aldrich), 2x Overlay Media (20% 10x MEM (Gibco), 2% L-glutamine (200mM) (Gibco), 2% Non-essential amino acids (Gibco), 6% Sodium bicarbonate solution (Gibco), 8% Fetal bovine serum (Sigma Aldrich), 5% HEPES buffer (Gibco) and 57% Distilled water (Versol). After 3 days incubation at 37°C, cells were fixed with formaldehyde and stained by addition of approximately 250μl 0.2% crystal violet for 5 minutes, before washing with water. The number of plaques in each well was determined and expressed as plaque forming units (pfu).

### RNA extraction and RT-PCR Analysis

RNA was extracted from aliquots (140 μL) of the coupon recovery liquid using the QIAamp Viral RNA Mini Kit (Qiagen Ltd, Manchester, UK). RT-PCR was performed using the VIASURE SARS-CoV-2 Real Time PCR Detection Kit (Viasure; CerTest Biotec, Zaragoza, Spain), following the methods provided. Quantification was undertaken using the N target with a standard curve generated by serial dilution of an in vitro transcript (10).

### Data Analysis

Each time point for each material coupon had 3 biological replicates (individual coupons) and 2 technical replicates (plaque assay performed in duplicate). Calculations for the mean and standard deviations of the assay counts were determined from these 6 replicates using Microsoft Excel (Office 365). Time to percent reduction values were calculated using linear regression of pfu/coupon averages, in Minitab 18. High and low loading line slopes were analysed using linear regression analysis with GraphPad Prism software version 7.

## Acknowledgements

Professor K Richards in High Containment Microbiology, PHE, Porton Down, for producing the viral stocks for this study.

The views expressed in this article are those of the authors and are not necessarily those of PHE or the Department of Health and Social Care. This manuscript is Crown copyright and is reproduced with the permission of Public Health England and the Controller of HMSO.

